# Impaired expected value computations coupled with overreliance on prediction error learning in schizophrenia

**DOI:** 10.1101/238089

**Authors:** D Hernaus, JM Gold, JA Waltz, MJ Frank

## Abstract

**Background:** While many have emphasized impaired reward prediction error (RPE) signaling in schizophrenia, multiple studies suggest that some decision-making deficits may arise from overreliance on RPE systems together with a compromised ability to represent expected value. Guided by computational frameworks, we formulated and tested two scenarios in which maladaptive representation of expected value should be most evident, thereby delineating conditions that may evoke decision-making impairments in schizophrenia.

**Methods:** In a modified reinforcement learning paradigm, 42 medicated people with schizophrenia (PSZ) and 36 healthy volunteers learned to select the most frequently rewarded option in a 75-25 pair: once when presented with more deterministic (90–10) and once when presented with more probabilistic (60–40) pairs. Novel and old combinations of choice options were presented in a subsequent transfer phase. Computational modeling was employed to elucidate contributions from RPE systems (“actor-critic”) and expected value (“Q-learning”).

**Results:** PSZ showed robust performance impairments with increasing value difference between two competing options, which strongly correlated with decreased contributions from expected value-based (“Q-learning”) learning. Moreover, a subtle yet consistent contextual choice bias for the “probabilistic” 75 option was present in PSZ, which could be accounted for by a context-dependent RPE in the “actor-critic”.

**Conclusions:** We provide evidence that decision-making impairments in schizophrenia increase monotonically with demands placed on expected value computations. A contextual choice bias is consistent with overreliance on RPE-based learning, which may signify a deficit secondary to the maladaptive representation of expected value. These results shed new light on conditions under which decisionmaking impairments may arise.

## Introduction

Reinforcement learning (RL) and decision-making impairments are a recurrent phenomenon in people with schizophrenia (PSZ) and are thought to play a key role in abnormal belief formation (1) and motivational deficits (2). While many have emphasized an impairment in learning from prediction errors (3–5) (the difference between expectation and outcome), multiple studies suggest that some of these deficits may in fact arise from overreliance on prediction errors together with a compromised ability to represent the prospective value of an action or choice (i.e. expected value) (e.g. (6, 7), for overview see Waltz & Gold (2)). However, such conclusions have typically been based on inferences, rather than experimental designs intended to reveal such effects. We therefore formulated and tested two hitherto unexplored scenarios motivated by the posited computations under which deficits in the representation of expected value should be most evident.

Optimal decision-making relies on a pas de deux between a flexible and precise representation of expected reward values, supported by orbitofrontal cortex (OFC) (8–10), which is complemented by a gradual build-up of stimulus-response associations, credited to dopaminergic teaching signals (reward prediction errors; RPEs) that project to striatum (11, 12). Previous work has demonstrated that maladaptive representations of expected value, rather than diminished stimulus-response learning per se, is one consistent feature of RL deficits in PSZ (13–15).

Findings of impaired representations of expected value in PSZ have often relied on computational models of learning and decision-making. In RL computational frameworks, it is thought that “Q-learning” and “actor-critic” models capture expected value and stimulus-response learning, respectively. In Q-learning (16), RPEs directly update the expected value of every choice option separately - similar to the representation of a reward value by OFC (17, 18) - and choices are driven by large action values. In contrast, in the actor-critic framework (19), choice preferences in the actor arise slowly on the basis of an accumulation of RPEs signaled by the critic, thought to reflect DA-mediated changes in synaptic weights in basal ganglia (20–22). Importantly, because the RPE fulfills different roles in these two computational frameworks (updating reward value directly versus modifying stimulus-response weights), it follows that, by definition, reward value is more precisely represented in Q-learning than in actor-critic frameworks. In one study, we showed that a computational modeling parameter that captured the balance between Q-learning versus actor-critic-type learning was tilted in favor of the latter in PSZ, suggesting relative underutilization of expected value and, perhaps secondarily, overreliance on stimulus-response learning (6). To date, however, little is known about the conditions under which deficits in the computation of expected value should be most observable.

We therefore sought to test two predictions of our theoretical account, which emphasizes maladaptive representation of expected value (Q-learning) in PSZ:

1. Counter-intuitively, and in contrast to many situations where PSZ may be most impaired at high levels of difficulty, our model based on less precise representations of reward value (decreased Q-learning) predicts that PSZ should suffer the largest decision-making deficits for the easiest value discriminations: that is, when the value difference between two competing options increases.
2. Secondly, if PSZ rely more on actor-critic-type learning - because of a decrease in Q-learning - then stimulus-response learning governs action-selection. In the actor-critic architecture, the tendency to repeat a choice or action is affected by the overall reward rate of the context, since RPEs are evaluated relative to that context. Therefore, a second diagnostic prediction is that context-dependent choice biases should be observable in PSZ even among items with identical reinforcement histories.

In the current study, we test these two hypothesized consequences of deficits in the representation of expected value using a modified RL paradigm. Participants were presented with two pairs of stimuli with identical reward value; one pair was presented in a “reward-rich” context (where the other pair had a higher reward rate), while the other pair was presented in a “reward-poor context” (where the other pair had a lower reward rate). Afterwards, participants were presented with old and novel combinations of choice options. We exploited the wide range in reward value to test our hypothesis relating to performance deficits as a function of the value difference between two competing options. Pairs with identical reward value in different contexts allowed us to address hypotheses relating to a contextual choice bias.

To accomplish these aims, we used a previously-validated hybrid computational model that estimates one’s tendency to use Q-learning *versus* actor-critic along a parametric continuum (6). As observed previously (6), we expected PSZ to rely less on Q-learning than actor-critic, resulting in the aforementioned deficits.

## Methods and Materials

### Sample

We recruited 44 participants with a Diagnostic and Statistical Manual of Mental Disorders, Version IV (DSM-IV) diagnosis of schizophrenia or schizoaffective disorder (PSZ) and 36 healthy volunteers (HV). Two PSZ were excluded; one participant was mistakenly administered an old version of the task, while another participant consistently performed below chance, leaving a sample of 42 PSZ. PSZ were recruited through clinics at the Maryland Psychiatric Research Center. A diagnosis of schizophrenia or schizoaffective disorder in PSZ, as well as the absence of a clinical disorder in HV, was confirmed using the SCID-I (23). The absence of an Axis II personality disorder in HV was confirmed using the SIDP-R (24). All PSZ were on a stable antipsychotic medication regimen. No changes in medication dose/type were made in the four weeks leading up to study participation. Major exclusion criteria included: pregnancy, current illegal drug use, substance dependence (in past year), a neurological disorder, and/or medical condition affecting study participation. All participants provided written informed consent. The study was approved by the Institutional Review Board of the University of Maryland SOM.

### Clinical ratings

We used the Scale for the Assessment of Negative Symptoms (SANS; (25)) to assess negative symptoms. The Brief Psychiatric Rating Scale (BPRS) positive symptom factor (suspiciousness, hallucinations, unusual thought content, grandiosity) was used as a measure of positive symptom severity (26). Antipsychotic doses were converted to haloperidol equivalents according to Andreasen et al. (27). All participants received the Wechsler Abbreviated Scale of Intelligence (WASI-II) (28) and the MATRICS Consensus Cognitive Battery (29). All ratings were collected by a trained and experienced clinical research associate.

### Reinforcement Learning Paradigm

Participants completed an RL paradigm consisting of a 320-trial learning phase and 112-trial transfer phase.

### Learning Phase

Participants were presented with pairs of stimuli and were asked to select one using their left (left stimulus) or right (right stimulus) index finger, after which they received positive (+$.05) or neutral ($0.00) feedback (figure 1). Choice feedback was delivered probabilistically according to three pre-determined contingencies (% positive feedback for optimal versus suboptimal choice): 1) 90-10, 2) 75-25, and 3) 60-40 (figure 1).

**Figure 1.**
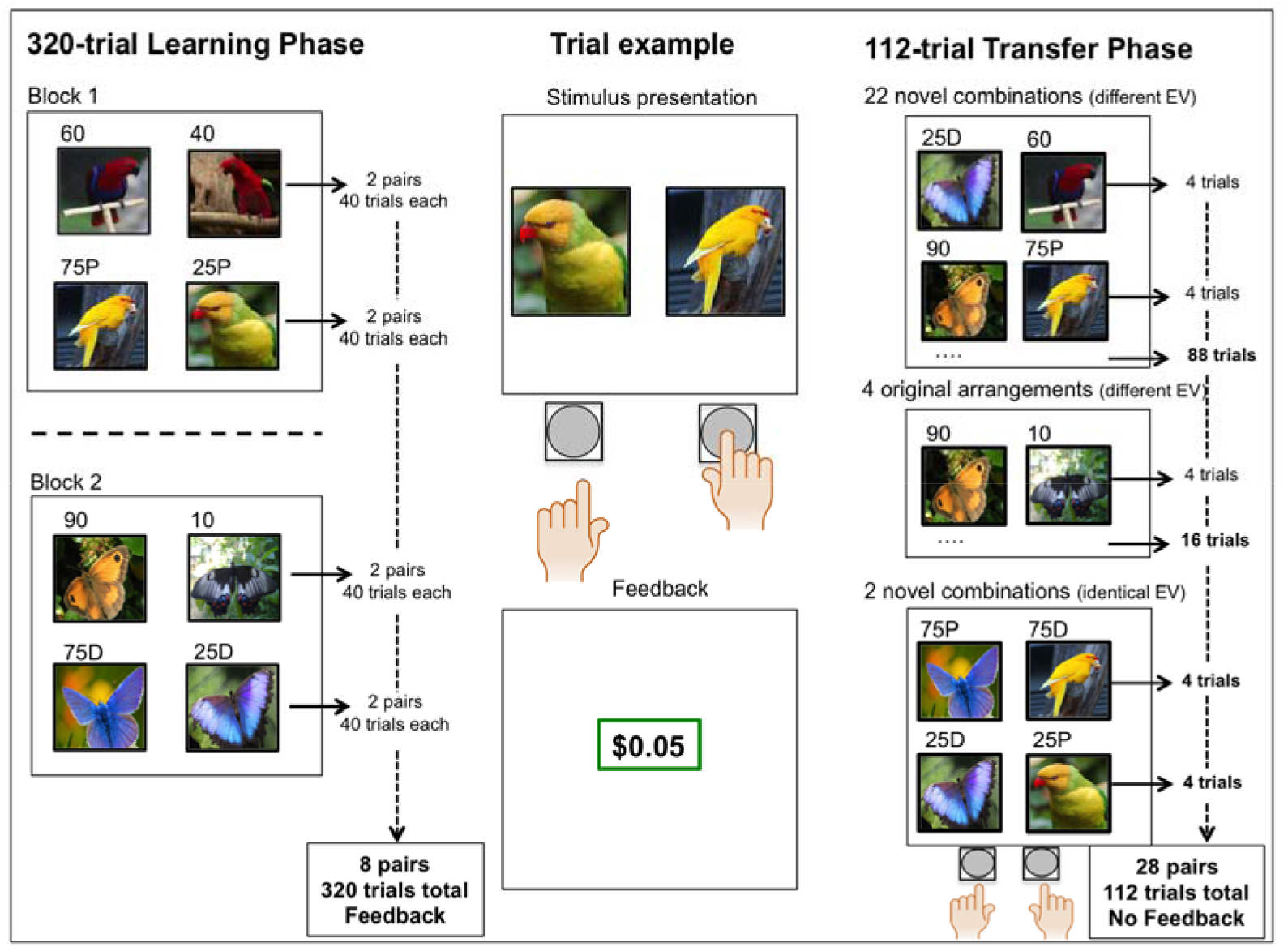
Overview of the reinforcement learning-and-transfer paradigm

The 320-trial learning phase was divided into two blocks of 160 trials. One block consisted of 80 90-10 and 80 75-25 trials, while the other block consisted of 80 75-25 and 80 60-40 trials. Trial presentation was pseudo-randomized within each block and block order was counterbalanced among participants.

In total, there were eight pairs; two 90-10 pairs, two 60-40 pairs, and four 75-25 pairs (every pair was presented 40 times). Stimulus type (butterflies/birds), option-probability pairing, and option-position pairing (left/right side of screen) were counter-balanced across participants. Note that two out of four 75-25 pairs always comprised bird-themed stimuli, and the other two pairs butterfly-themed stimuli (figure 1).

By combining 75-25 pairs with more deterministic (90–10) and probabilistic (60–40) pairs in separate blocks, we aimed to investigate context-dependent RL, meaning that perceived choice value (here, 75-25 pairs) might be dependent on contextual reward rate (the average reward rate of optimal choice options within a block). Henceforth, we will refer to 75-25 pairs that were presented together with 90-10 pairs as “75-25D” (“D” for the more deterministic context) and 75-25 pairs that were presented with 60-40 pairs as “75-25P” (“P” for the more probabilistic context).

### Transfer Phase

The 112-trial transfer phase served two purposes: I) to assess the ability to compare choice options using their reward value, and, II) a formal test of a contextual choice bias.

Every possible combination of two choice options (new and original combinations) was presented and the participant was instructed to “select the option that was rewarded most often” (figure 1). To prevent further learning, no feedback was delivered. Combining all possible choice options yielded twenty-eight combinations with non-identical expected value; twenty-two novel combinations (e.g. 90-60 or 75D-10) and four original combinations (90-10, 75-25D, 75-25P, 60-40). In addition, we produced two novel combinations with identical expected value; 75P-75D and 25P-25D. Supplementary table 1 gives an overview of transfer pairs that were used for the analyses described below.

Note that there were two pairs of each contingency in the learning phase, with the exception of 75-25 pairs, of which there four. Thus, although every unique combination of choice options was presented only once in the transfer phase, there were always four presentations of each expected value combination (e.g. with two 90-10 pairs one can generate four unique 90-10 combinations).

### Computational model Hybrid Model

In an attempt to relate deficits in expected value and a contextual choice bias to latent variables, we used a previously-validated hybrid model allowing for combined influences of Q-learning and actor-critic frameworks on decision-making (6). The model with the best trade-off between model complexity, fit, and posterior predictive simulations contained six free parameters: a critic learning rate (αc), actor learning rate (α_a_), Q learning rate (α_q_), beta (β; inverse temperature), mixing (m), and undirected noise (ε) parameter, which were estimated for every subject via maximum likelihood optimization (see supplemental text and supplemental figure 1 for a detailed description of the model selection procedure).

In actor-critic models, the critic learns state values, while the actor selects responses (19, 30). The critic learns about state value by observing outcomes and uses its own RPE [δ(t) = reward r(t) – expectation q(t)] to update the state value of a stimulus pair [state value v(t) = v(t-1) + α_c_*δ(t)]. The critic’s RPE also updates the actor’s action weights for chosen options [action weight aw(t) = aw(t-1) + α_a_*δ(t)]. Thus, action-weights - similar to action values in Q-learning - shape choice preferences. Collins and Frank (21) showed that a modified actor-critic was able capture the contributions of basal ganglia to a wide array of data.

In contrast, in Q-learning models, RPEs are used to directly estimate an option’s expected value [action value av(t) = av(t-1) + α_q_*δ(t)], which governs future choices (16, 17, 19). Thus, there is an important conceptual distinction between actor-critic and Q-learning; whereas the former can develop a response tendency without knowing the value of a choice (because the actor does not observe outcomes and is informed only by the critic), the latter develops a value representation for every choice. A precise representation of action value, as in the Q-learning model, might be especially relevant when presented with new combinations of old choice options (e.g. 90>10, 60>40, thus 90>60). The m parameter in our hybrid model allows us to estimate parametric mixing between these two frameworks; an m parameter <.5 suggests greater reliance on actor-critic learning, whereas an m parameter >.5 suggest a greater contribution of Q-learning. Lastly, ε accounts for undirected noise, with more random behavior as ε approaches 1. Individual estimates of ε were used in the softmax function to allow for randomness in decision-making unrelated to learned values/weights (see Nassar and Frank (31) and references therein).

### Contextual choice bias

The critic in actor-critic models represents the value of the overall state (19), where dopamine signals are thought to reflect RPEs relative to that state. In light of reward rate in the generation of RPEs (32), we estimated a single state value (V) for each block, similar to “average reward” RL, where rewards are interpreted in relation to the long-term reward rate (33). Context-dependent state values allow for the possibility that individuals learn stronger response tendencies (actor-weights) toward 75% rewarding options learned in a context with relatively low reward rate (which should produce a greater RPE relative to the context) than those for 75% stimuli in contexts with high reward rate. Importantly, however, a contextual choice bias would only be present to the degree that participants relied on actor-critic type learning. If participants relied solely on action values, as in Q-learning, then contextual reward availability would not affect choice preferences.

After fitting the hybrid model to the learning phase data, the final action weights of all eight original pairs were used to simulate transfer phase performance for all pairs (n(simulations)=250 for every participant).

### Statistical analyses

Learning phase performance on every reinforcement contingency was averaged (per two pairs) and grouped into four bins of 20 trials. A 2x4x4 repeated-measures ANOVA using group status (predictor) and reinforcement contingency (4 levels) and trial-bin (bins; 4 levels) as dependent variables was run to test for a group-by-condition-by-time interaction. Group-by-time and group-by-condition interactions were also investigated. Greenhouse-Geisser sphericity-corrected values were reported when assumptions were violated.

Transfer phase accuracy was averaged across all four presentations of every unique (28) combination of expected values and compared using two-sample t-tests. Transfer phase pairs were next ranked on their value difference (supplementary table 1 for details regarding these outcome measures). A logistic regression analysis with value difference (left-right option) as predictor and correct choice (left vs. right button) as dependent variable was conducted to test the hypothesis that PSZ show impaired performance with increasing value difference. Individual value difference slopes were compared in a two-sample t-test.

Context-dependent learning (75P-75D trials) was investigated using a two-sample t-test, as well as a one-sample t-test to compare preference for either option against chance. As an indirect measure of context-dependent learning, performance on all trials where 75P and 75D stimuli were presented with any other option (excluding the 25 stimulus that they were originally partnered with) was compared in a 2x2 group-by-pair ANOVA Correlation analyses with clinical and psychometric variables were carried out using Pearson’s r and Spearman’s ρ (when distributions were skewed). Significance thresholds were set to p<.05.

## Results

### Demographics

Participant groups were matched on most demographics. However, PSZ did have a lower IQ-score than HV, as well as poorer MATRICS performance (table 1).

**Table 1.**
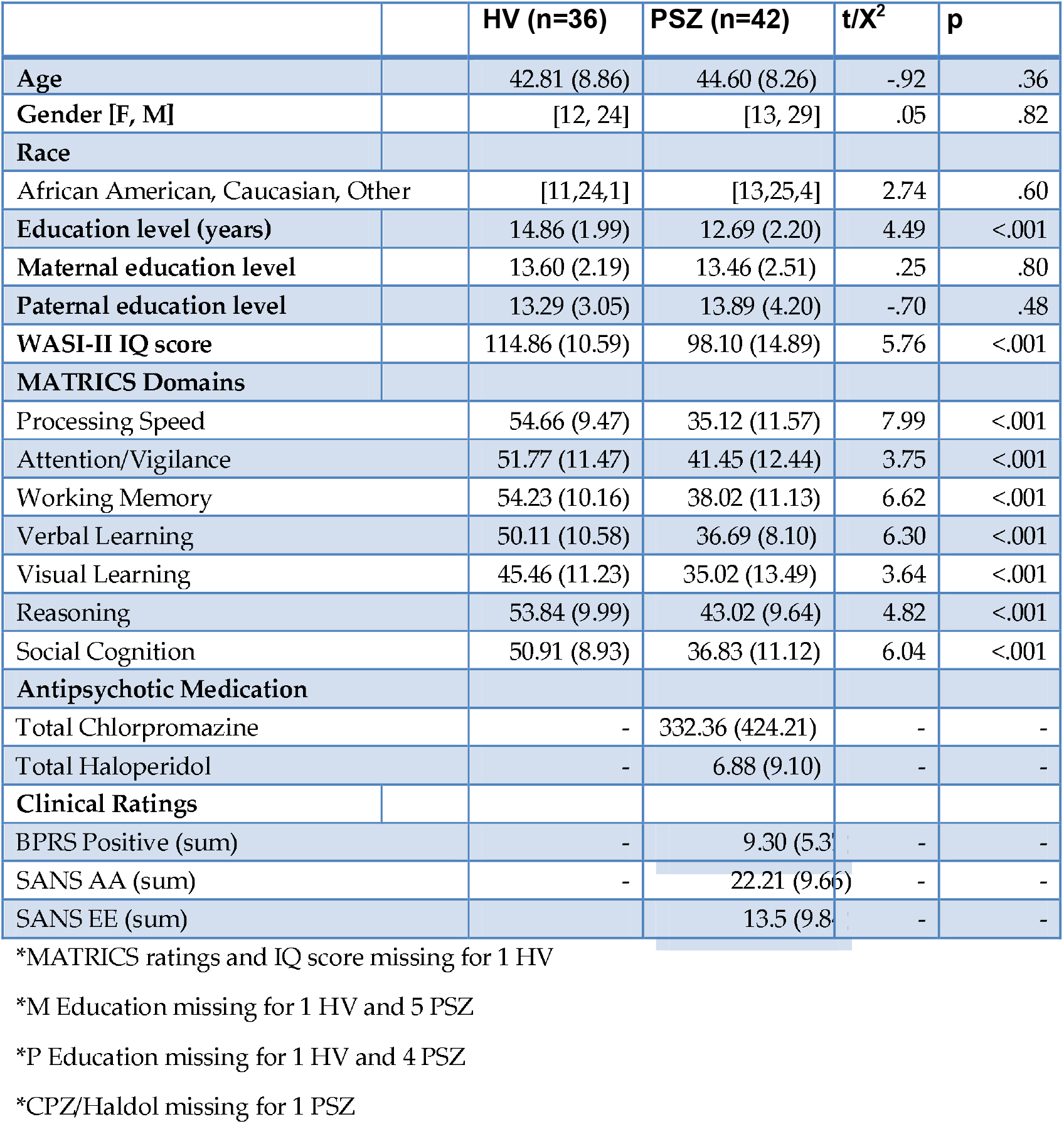
Demographics

### Learning Phase performance

We observed a group*probability interaction (F_3,228_=4.39, *p*=.005), such that HV outperformed PSZ in the 90-10 (*p*=.002), 75-25D (*p*=.007), 75-25P (*p*=.04), but not 60-40 (*p*=.63), probability condition (Figure 2A). Group*probabi I ity*ti me (F_9,684_=.98, *p*=.46) and group*time (F_3,228_=.62, *p*=.60) interactions were not significant. Performance on 60-40 trials in bin 4 was significantly above chance for both groups (HV: t_35_=2.88, *p*=.007; SZ: t_41_=2,64, *p*=.01)

**Figure 2.**
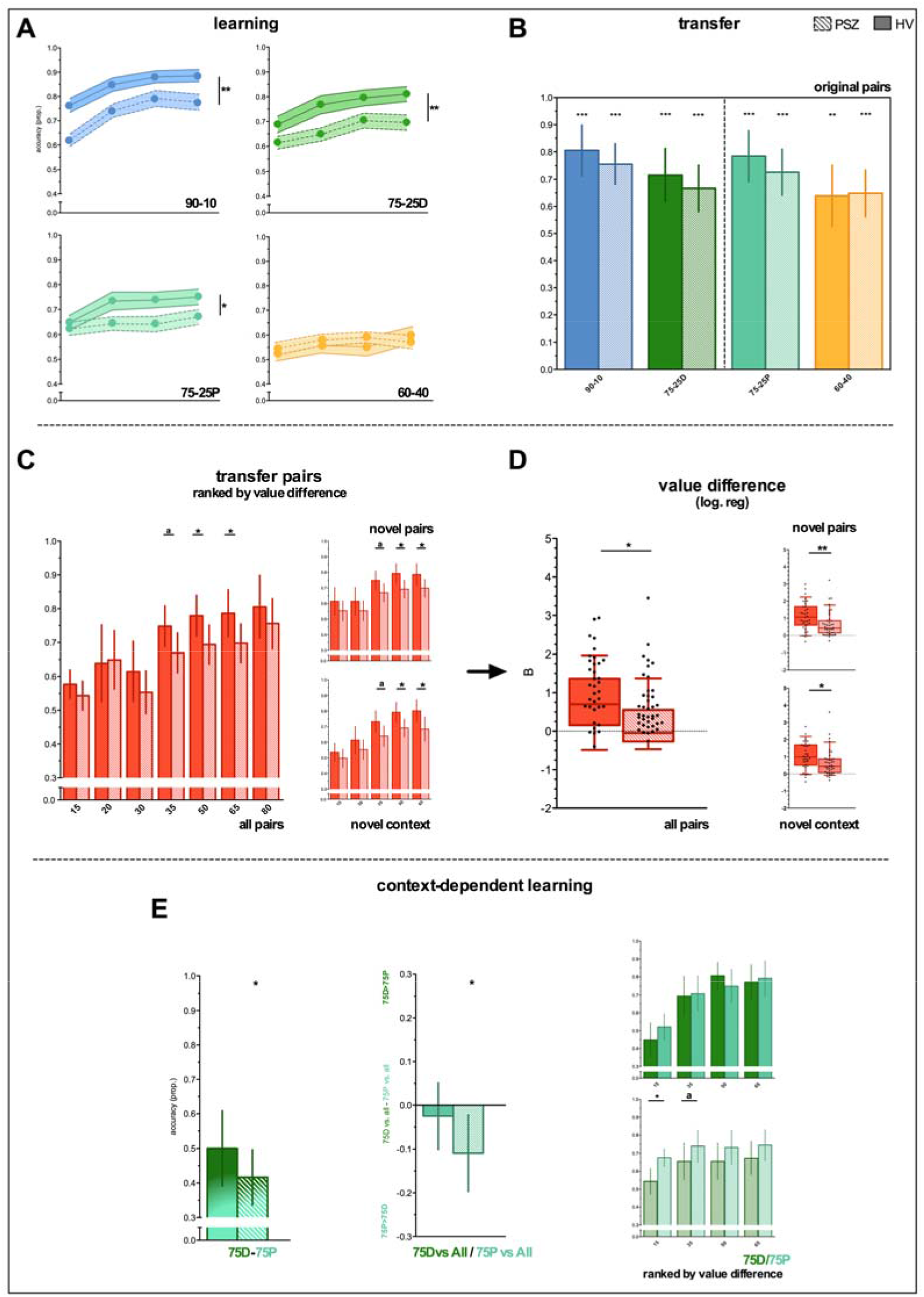
Learning and transfer phase performance. Solid bars represent HV; bars with diagonal lines represent PSZ. *= p<.05, ** = p<.01, = p<.001, a = trend (p = .06-.09). Error bars represent 95% CI, except for learning phase data, where bars represent SEM. Asterisks above error bars represent significant preference against chance; asterisks above solid horizontal line represent between- or within-group differences. Bottom row, center: “75D vs. All/75 vs. All” = performance on 75D/P trials *versus* all choice options of non-identical value. Bottom row, right: separate plots for 75D and 75P *versus* other choice options broken down by their value difference (x-axis).

### Transfer Phase performance

Despite poorer learning accuracy in PSZ there were no group differences in transfer accuracy for 90-10, 75-25D, 75-25P or 60-40 pairings (all *p*>.39; Figure 2B), with accuracy above chance on all pairs.

### Smaller performance improvements with increasing value difference in PSZ

Accuracy on all novel pairs is shown in Supplemental Figure 2. When all combinations of reward contingencies were considered, accuracy on trials with a value difference of 35 (t_77_=3.55, *p*=.06), 50 (t_77_=4.26, *p*=.04), and 60 (t_77_=4.08, *p*=.05) were (trend-wise) greater in HV compared to PSZ (Figure 2C). This was also true when using only novel pairs or only pairs consisting of one choice option from each context (Figure 2C). To formally test the presence of a greater accuracy deficit with increasing value difference, we compared individual slopes from a logistic regression predicting accuracy as a function of value difference. Using all pairs (t_74_=5.84, *p*=.02), novel pairs (t_74_=6.99, *p*=.01), and novel context pairs (t_73_=6.05, *p*=.02), the slope for HV was always greater than PSZ (these results could not be used in 4 participants due to limited choice variability; Figure 2D). This all suggests that PSZ, compared to HV, improved less as the value difference between two competing stimuli increased, thereby confirming our first initial hypothesis.

### Context influences perceived stimulus value in PSZ, but not HV

A direct comparison of 75D-75P performance revealed no significant group difference (t_77_=1.61, *p*=.21) (Figure 2E). PSZ (one sample t-test against chance: t_41_= -2.10, *p*=.04), but not HV (t_41_=0.01, *p*=.99), did however show a significant preference for 75P over 75D. The more indirect group*pair interaction for 75P and 75D performance *versus* other options showed similar numerical patterns but was not significant (t_1,67_=2.11, *p*=. 15). Nevertheless, PSZ (t_41_=-2.52, *p*=.015), but not HV (t_35_=-.67, *p*=.51), more often selected 75P than 75D when paired with another option (Figure 2E). The direct and indirect measure of context-sensitivity correlated in PSZ (Pearon’s *r*= -.53, p<.001) Taken together, these results provide subtle yet consistent evidence that context may impact perceived choice value in PSZ, but not HV. To formally test whether the trial-by-trial pattern of choices can be explained by context-dependent value learning, we next turn to computational model results.

### Computational modeling results

As predicted, the mixing (*m*) parameter was significantly greater in HV than PSZ (t_76_=2.51, *p*=.01), suggesting that PSZ relied more on actor-critic, and less on Q-learning, than HV. In addition, the undirected noise parameter was greater in PSZ than HV (t_76_=2.52, *p*=.01) (Figure 3A; supplementary table 2 for parameters per subject).

**Figure 3.**
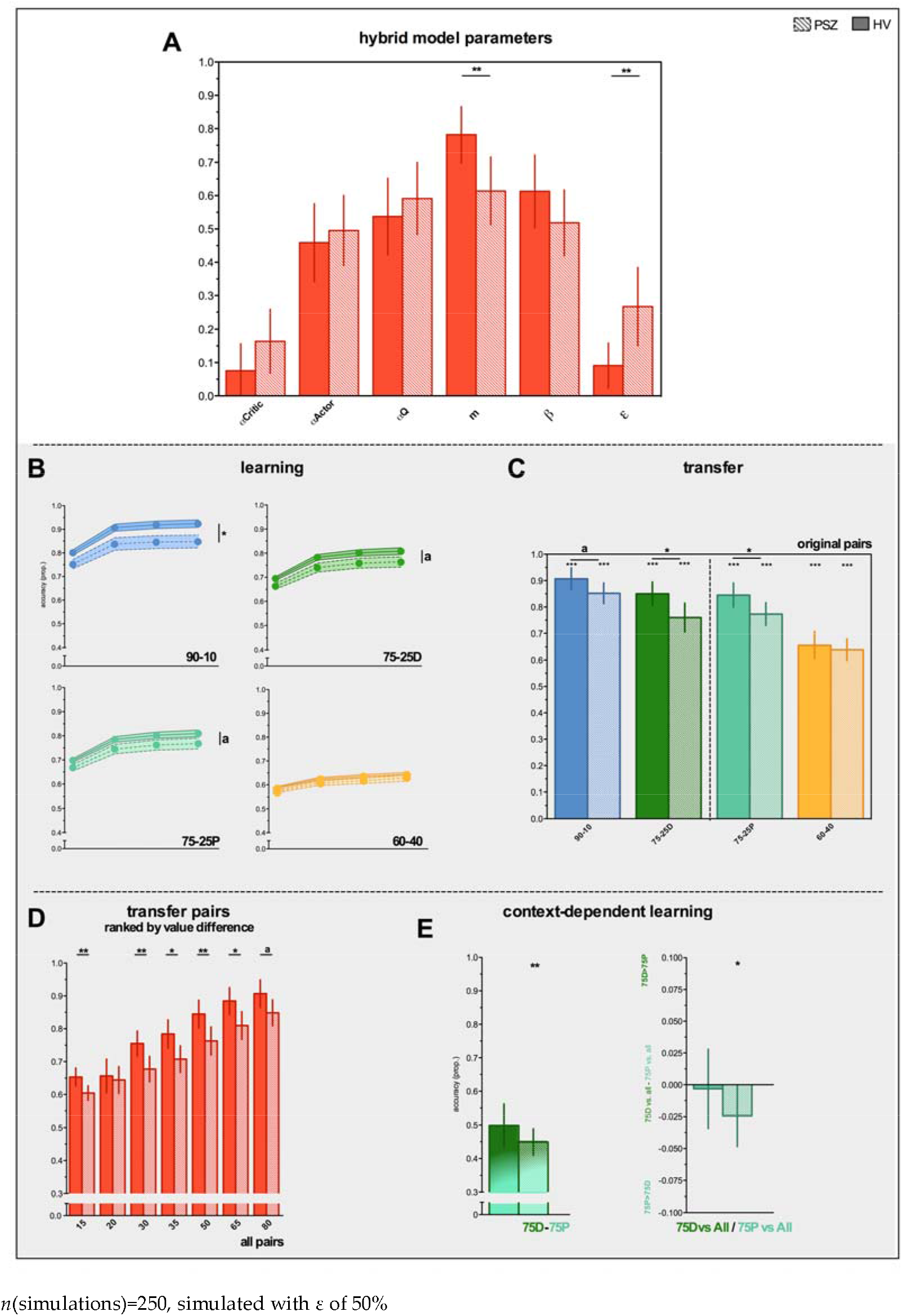
Hybrid model parameters and simulated data *n*(simulations)=250, simulated with ε of 50%

True to the actual learning phase data, model simulations revealed numerically greater performance in HV relative to PSZ for 90-10, 75-25D, 75-25P (but not 60-40) contingencies, which became (trend-)significant when increasing the number of simulations (*n*(simulations) = 1000 shown in Figure 3B). Because the number of trials in the transfer phase was relatively small (four per combination) and the amount of undirected noise may be greater during learning compared to transfer phase performance, we set ε to 50% of the original value during transfer phase simulations.

All findings remained when simulating transfer data with ε set to 100% (Supplemental Figure 3).

Predicted group differences in transfer phase accuracy on 90-10, 75-25D, and 75-25P pairs were small, as was the case in the original data, yet (trend) significant, owing to the number of simulations (Figure 3C; w(simulations) for all transfer data = 250). Importantly, the hybrid model predicted numerically greater performance deficits in PSZ with increasing value difference (Figure 3D). The direct and indirect context effects in PSZ were both present in the simulated data (Figure 3E): that is, I) a preference for 75P over 75D (t_40_=2.59, *p*=.01) and II) a preference for 75P over 75D when paired with all other stimuli (t_39_=2.03, *p*=.05). One outlier in the PSZ sample with high values overall/difference scores was removed from the simulated data; excluding this subject from the actual data did not change the results.

### Correlations among performance measures, model parameters and clinical variables

The *m* (Spearman’s ϱ=-.67, *p*<.001) and ε (Spearman’s ϱ=.38, *p*<.001) parameter significantly correlated with the slope of the value difference effect in the entire sample, suggesting that decreased reliance on Q-learning and greater undirected noise were associated with smaller performance improvements with increasing value difference (Figure 4A/4B).

**Figure 4.**
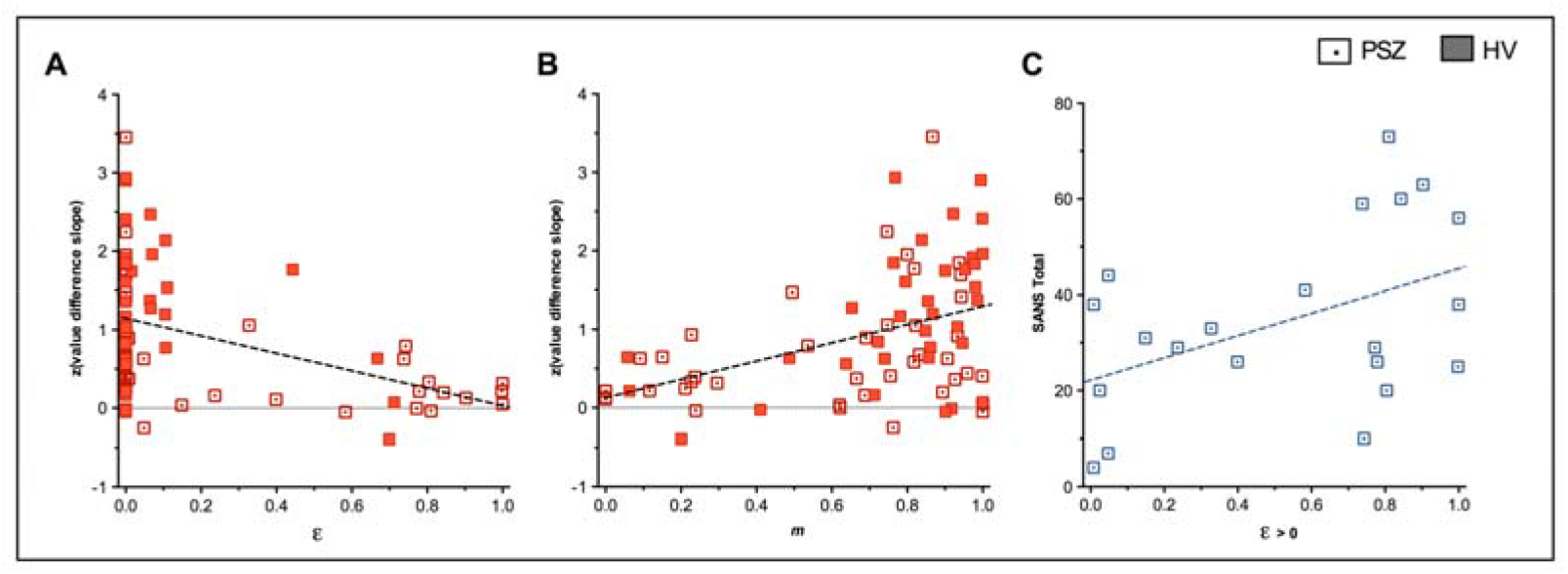
Significant correlations among model parameters, symptoms, and performance

Next, we focused on the α_c_ parameter, which can produce the context effect within the actor-critic model. To demonstrate this, *m* and ε were both fixed at 0 and the direct and indirect context effect were simulated, thereby removing contributions from Q-learning and undirected noise, while all other parameters were set to original values. In PSZ, α_c_ correlated with the size of the simulated direct (Spearman’s ϱ=-.42, *p*<.005) and indirect (Spearman’s ϱ r=.47, *p*<.001) context effect. This confirms our intuition that varying levels of critic learning rate are sufficient to account for the context effect. Moreover, when simulating data using individual *m* and ε parameters, α_c_-weighted [α_c_*(l-m)] also significantly correlated with the simulated indirect context effect (Pearson’s r=.34, *p*=.02), while the correlation with the simulated (Pearson’s r=-.28, *p*=.07) and actual (Pearson’s r=-.13, *p*=.41) direct context effect was in the expected direction but not significant. This provides evidence that greater α_c_ values can account for a context-dependent choice bias, although this also crucially depends on the degree to which participants rely on Q-learning and the amount of undirected noise.

Finally, we looked at associations with clinical ratings. In PSZ with some degree of randomness (i.e. ε>0; n=21), there was a trend for ε to correlate positively with SANS total scores (Spearman’s ϱ=.42, p=.06), suggesting that individuals with greater motivational deficits may show more undirected noise. SANS total scores did not correlate with m (Pearson’s r=.14, p=.38). M also did not correlate with IQ (HV Spearman’s ϱ=.28 p=.10: SZ Spearman’s ϱ=.18, p=.27). In PSZ (Spearman’s ϱ=-.36, p=.02) and (trend-wise in) HV (Spearman’s ϱ=.31, p.06), ε correlated with IQ. Finally, at correlated with IQ in PSZ (Spearman’s ϱ=-.34, p=.03), but not in HV (Spearman’s ϱ=-.18 p=.31).

M, ε and α_c_ did not correlate with haloperidol equivalents (all p>.71), BPRS positive symptoms (p>.31) or age (p>.32).

## Discussion

Using theory-based predictions, our primary aim was to investigate two hypothesized RL and decision-making deficits that could result from a relative underutilization of expected value. As predicted, PSZ showed robust performance impairments as the difference in reward value between two choice options increased. Moreover, we observed a subtle yet consistent contextual choice bias that was not present in controls: when presented with two options of identical reward value (75D and 75P), or when these options were paired with options of other reward value, PSZ preferred the 75 option from the more probabilistic context (75P).

Performance deficits amplified at greater levels of value difference are diagnostic of a change in the choice function rather than a general learning impairment, which would typically manifest in the opposite manner: that is, worse performance for more difficult judgments. These results are particularly noteworthy because they further corroborate the notion that certain learning and decisionmaking deficits in PSZ are associated with a selective deficit in the representation of expected value. A more general learning impairment, potentially via altered RPE signaling of midbrain DA neurons (1, 3, 5), would predict that performance impairments in PSZ increase in conditions where the value difference between two choice options is subtle. We have previously observed a hint for performance deficits at greater levels of value difference in other RL tasks (6, 34), suggesting that this is a recurrent impairment in PSZ. Our computational model provides evidence that such impairments stem from a decrease in action value (Q-) learning (via the m parameter). Importantly, these results conceptually replicate our previous work for the first time, in which we showed a decreased contribution of Q-learning during a gain-seeking/loss-avoidance task (6). In the current study, performance impairments were also in part related to increased undirected noise (ε), which accounts for non-deterministic choices even in the face of strong evidence. We have observed this in previous reinforcement learning studies (15), and, in the current study, was mostly driven by PSZ with greater motivational deficits.

Which mechanisms could underlie a selective impairment in the representation of expected value? Decreased learning from gains, as opposed to intact loss-a voidancc, has been identified as one potential mechanism (6, 14, 35, 36). In this study, impaired performance on more deterministic pairs, associated with more gains than neutral outcomes, but spared performance on 60-40 trials, where learning occurs almost equally from gains and neutral outcomes, provides circumstantial evidence for this notion. One improvement compared to previous paradigms is that, here, we focused on reward value instead of contrasting valence conditions, which is a more direct test of expected value deficits. The current results show for the first time that a diminished role of expected value in driving choices can lead to suboptimal behavior in a dose-response fashion; that is, performance impairments increase monotonically with increased demands placed on expected value computations. This work further strengthens the claim that deficits in the representation of expected value are a central feature of learning and decisionmaking impairments in schizophrenia, and here we reveal when these deficits should be most evident.

The relationship between the value difference effect and Q-learning fits well with previous neuroimaging studies. Previous work has identified attenuated expected value signals in insula and anterior cingulate, regions that encode (state-dependent) expected value (37, 38), in PSZ with motivational deficits (5, 14). Ventromedial and orbitofrontal prefrontal cortex dysfunction, consistently involved in tracking reward value (8, 9, 39), has also been linked to learning and decisionmaking deficits in schizophrenia (40, 41). Thus, a diminished role for expected value in decision-making, demonstrated by the value difference effect and confirmed by our computational model, are suggestive of impairments in a range of cortical areas that encode reward value.

We have argued that underutilization of expected value and increased reliance on stimulus-response learning can also enhance the effect of context on stimulus valuation, leading to a unique prediction in which preferences can arise among choice options with identical reinforcement probabilities. For this hypothesis we found subtle but consistent evidence in PSZ, but not controls, which could be accounted for by a context-dependent state-value RPE (via α_c_). Although the effect of contextual reward availability on decision-making was subtle in PSZ, these findings are noteworthy. Klein et al (42) revealed that learning the value of one stimulus relative to another can lead to sub-optimal decision-making. In their study, a relative RPE signal was specifically encoded by the striatum. Despite clear differences between the task design of Klein et al. (42) and the current study, most notably pairwise versus block-wise context effects, their work does provide evidence for the notion that the effect of context on perceived stimulus value seems to be encoded specifically by brain regions typically associated with RPE signaling.

Related to this point, we did not find evidence of group differences on 60-40 trials (also see Waltz et al. (34)), where performance improved gradually and likely relies on slow accumulation of RPEs (17). Subtle evidence for a context effect, a relative increase in the contribution of actor-critic-type learning, and no group difference in performance on 60-40 trials are consistent with relatively intact striatal function in our medicated sample. These findings align well with intact striatal RPE signaling in medicated individuals with schizophrenia (43) as well as normalization of reward signals following treatment with antipsychotics (44). Given evidence of abnormal RPE signals in unmedicated individuals with psychosis (45), studies into a contextual choice bias in unmedicated participants could promote further understanding of the degree to which striatal stimulus-response learning might be involved.

To summarize, this work provides specific evidence that decision-making impairments in schizophrenia increase monotonically with demands placed on expected value computations. Overreliance on stimulus-response learning as a result of underutilization of expected value may produce additional violations of optimal decision-making policies, such as a contextual or relative choice bias. This work provides a novel source of evidence suggesting a diminished role of expected value in guiding optimal decisions in schizophrenia and sheds light on the conditions that facilitate such impairments.

### Limitations

Some limitations warrant discussion. While we were able to replicate our previous finding of decreased Q-learning/increased actor-critic learning in PSZ (6), the m parameter was not associated with symptom ratings. Previous studies investigating RL deficits in schizophrenia have reported mixed results regarding relationships to negative symptoms (6, 35, 46). Clinical status, symptom severity, and experiment design (notably, the wide range in reinforcement contingencies and the emphasis on expected value computations) might explain some of these discrepancies.

Moreover, an alternative account of the current findings is that PSZ may rely less on model-based strategies (47). Both Q- and model-based learning make identical predictions for this task: that is, Q-learning predicts improved performance at greater levels of value difference via action-value learning, while model-based strategies predict improved performance when action-outcome sequences are better understood. Importantly, this alternative explanation does not change the interpretation of increased reliance on model-free stimulus-response learning in PSZ.

## Acknowledgements

We thank Benjamin M Robinson for his contributions to the task design.

